# Genome-skimming provides accurate quantification for pollen mixtures Authors

**DOI:** 10.1101/408039

**Authors:** Dandan Lang, Min Tang, Jiahui Hu, Xin Zhou

## Abstract

In the face of global pollinator declines, plant-pollinator interaction networks have been studied to guide ecological conservation and restoration. In order to obtain more comprehensive and unbiased knowledge of these networks, perspectives of both plants and pollinators need to be considered integratively. Metabarcoding has seen increasing applications in characterizing pollen transported by pollinators. However, amplification bias across taxa could lead to unpredictable artefacts in pollen compositions. We examined the efficacy of a PCR-free genome-skimming method in quantifying mixed pollen, using mock samples constructed with known pollen species (5 mocks of flower pollen and 14 mocks of bee pollen). The results demonstrated a high level of repeatability and accuracy in identifying pollen from mixtures of varied species ratios. All pollen species were detected in all mock samples, and pollen frequencies estimated from the number of sequence reads of each species were significantly correlated with pollen count proportions (linear model, R^2^ =86.7%, P = 2.2e- 16). For >97% of the mixed taxa, pollen proportion could be quantified by sequencing to the correct order of magnitude, even for species which constituted only 0.2% of the total pollen. We also showed that DNA extracted from pollen grains equivalent to those collected from a single honeybee corbicula was sufficient for the genome-skimming pipeline. We conclude that genome-skimming is a feasible approach to identifying and quantifying pollen compositions for mixed pollen samples. By providing reliable and sensitive taxon identification and relative abundance, this method is expected to improve the understanding of pollen diversity transported by pollinators and their ecological roles in the plant-pollinator networks.

## Introduction

Pollinator declines have been widely reported in the last decades, causing substantial losses in pollination services and subsequent reductions in crop yields (Potts et al., 2010). Over 80% of known flowering plants are pollinated by animals, mainly insects (Ollerton, Winfree, & Tarrant, 2011). Therefore, global conservation efforts have been carried out with a priority on biodiversity registration and monitoring of pollinators, especially bees (Winfree, Griswold, & Kremen, 2007; Burkle, Marlin, & Knight, 2013). In addition, an increasing amount of studies have been focused on the understanding of pollination functions of pollinators through construction of pollination networks (Kremen, 2005; McCann, 2007; Tylianakis, Laliberté, Nielsen, & Bascompte, 2010; Schleuning, Fründ, & García, 2014). In turn, the network and its resilience to disturbances have been suggested as an effective proxy for evaluating the efficacy of conservation management and restoration of ecosystem function (Devoto, Bailey, Craze, & Memmott, 2012; Kaiserbunbury & Blüthgen, 2015; Kaiser-Bunbury et al., 2017).

Plant-pollinator network studies often involve field observation of flower visitations by various pollinators. However, not all flower visitors provide pollination service (King, Ballantyne, & Willmer, 2013) and when they do, they may vary in efficiencies in pollen transport and pollination success (Memmott, 1999). In addition, the construction of visitation networks is typically time-consuming (e.g., hours of observation on a moderate network containing a few dozens of flower species, Kaiser-Bunbury et al., 2017), which usually leads to under-sampling of flower diversity or replicates, especially when working at large geographical scales. Unfortunately, such under-samplings tend to underestimate the level of nestedness (the degree to which interactions of specialist species are residing in that of more generalist species) (Petanidou, Kallimanis, Tzanopoulos, Sgardelis, & Pantis, 2008; Bosch, González, Rodrigo, & Navarro, 2009), a crucial feature indicating resilience of the network to perturbations (Bascompte, Jordano, Melián, & Olesen, 2003; Rohr, Saavedra, & Bascompte, 2014).

As a complementary approach, pollen transport web based on analysis of pollen loads carried by insects (Forup & Memmott, 2005) may help to fine-tune pollination networks constructed from visitations (Kanstrup & Olesen, 2000; Gibson, Nelson, Hopkins, Hamlett, & Memmott, 2006; Forup, Henson, Craze, & Memmott, 2008; Jedrzejewska-Szmek, Krystyna, Zych, & Marcin, 2013). Firstly, pollen analysis of flower visitors would help characterize the diversity of pollen carried by animals, therefore identifying pollen transport variations, through which visitors carrying no pollen could be excluded from the network. Secondly, pollen loads provide extended visitation information for visitors (over time and space) as pollen grains would stay on pollinators’ bodies when they travel across flowers (Courtney, Hill, & Westerman, 1982). In fact, pollen analysis have been demonstrated to complement network interactions built from visitations by increasing pollinator connectivity, nestedness and centralization, and by revealing new modules (Bosch et al., 2009).

However, pollen identification based on classic palynology requires specialized skill set. Pollen grains are usually stained and identified morphologically under a microscope, which is time- and labor-consuming. Furthermore, rare species are prone to be overlooked in subsampling and microscopic examination. Alternatively, molecular identifications especially metabarcoding have seen increasing applications in bulk pollen characterizations (Galimberti et al., 2014; Richardson et al., 2015; Cornman, Otto, Iwanowicz, & Pettis, 2015; Keller et al., 2015; Bell et al., 2016; Danner, Molitor, Schiele, Härtel, & Steffan-Dewenter, 2016; Pornon et al., 2016; Bell et al., 2017; Kamo et al., 2018). Metabarcoding employs high-throughput sequencing (HTS) in analyzing pooled amplicons obtained from mixed taxa (Ji et al., 2013; Cristescu, 2014). While PCR of target genes (e.g., DNA barcodes) helps to increase DNA quantity for HTS, this procedure is prone to introducing taxonomic bias due to varied primer efficiencies across taxon lineages (Crampton-Platt, Yu, Zhou, & Vogler, 2016), and multiple optimization methods have been proposed (e.g., Nichols et al., 2018; Piñol, Senar, & Symondson, 2018). Recent studies have introduced a PCR-free approach, a.k.a. genome-skimming, where the total DNA extracts from bulk samples are directly subject to shotgun sequencing, therefore providing better qualitative and quantitative results for pooled invertebrate samples (Zhou et al., 2013; Tang et al., 2015; Arribas, Andújar, Hopkins, Shepherd, & Vogler, 2016; Choo, Crampton-Platt, & Vogler, 2017; Bista et al., 2018).

Quantitative composition of pollen mixtures is particularly important to the understanding of the flower diversity that bees visit and potentially their pollination contribution to the ecosystem. However, the feasibility of genome-skimming in pollen samples has not been proofed. In particular, as pollen grains are typically small in sizes, they may not provide sufficient DNA for HTS without PCR amplifications. In this study, we examined the potentials of pollen genome-skimming using mock samples consisting of known pollen at varied ratios. We show that our approach can provide accurate taxon identification for pollen mixtures and quantitative information for all member species, including those presented at low abundances. We also demonstrate that our method is feasible with small amount of pollen, where pollen pellets carried by individual bees can provide sufficient DNA for genome-skimming. Finally, we discuss practical considerations in the incorporation of new method into current pollen network studies, including analytical cost, operational complexity and compatibility with existing methods.

## Materials and Methods

### Pollen Samples

Flower pollen (FP) was collected from fresh flowers (*Abutilon megapotamicum*, *Ab. pictum*, *Alstroemeria aurea*, *Antirrhinum majus*, *Lilium brownii*, *Nymphaea stellate* and *Schlumbergera truncates*), which were purchased from a local flower market. Fresh flowers were identified morphologically by Dr. Lei Gu of Capital Normal University, China. Mature pollen grains were sampled with a sterile needle and preserved in a sterile vial for each species. Before pollen maturation, stamens from each species were isolated in petri dishes separately to avoid cross-contamination. Bee pollen (BP) of *Brassica napus, Camellia japonica, Papaver rhoeas, Prunus armeniaca, Rhus chinensis* and *Vicia faba* were purchased from Internet stores. These pollen pellets were collected from corbiculae (pollen baskets) of farm honeybees (*Apis mellifera*), then desilicated and bottled by the merchandise. Pollen identity and composition were examined using DNA barcoding and shotgun sequencing (described in the following paragraphs). Two grams of each BP (ca. 200-350 pollen pellets) were dissolved with sterile water then centrifuged at 14,000 g for 10 minutes and the supernatant was gently removed. FP and BP were suspended with 1 mL and 20 mL of 95% ethanol, respectively.

### Pollen Counting

Subsamples of FP and BP pollen suspensions were added into the Fuchsin dilution (16% glycerol, 33% alcohol, 1% basic fuchsine dye, and 50% deionized water) for pollen counting, where the total volumes were adjusted so that individual pollen grains could be recognized under a microscope without overlapping (Table S1 in Appendix 1). Pollen Fuchsin suspensions were homogenized by vortex shaking and a 5 μL or 10 μL subsample was then examined on a glass slide under a Nikon SMZ800N microscope or on a blood cell counting plate under a Nikon SMZ745T microscope, for FP and BP, respectively (almost all BP had smaller grains than FP in the studied species). To reduce stochastic errors during the process, the dilution procedure for each pollen species was repeated 3 times, whereas counting was repeated 3 times for each dilution. The average count from these 9 replicates was considered as the final pollen count for that species. These counts were then used to calculate pollen numbers per volume unit for each species.

### Pollen Mixture Mocks

Mock samples with species mixed at varied proportions each contained 200,000 to 5,000,000 pollen grains (Table 1), roughly reflecting those carried by an individual honeybee on 1 or 2 corbiculate legs (estimated by BP samples, Table S1 in Appendix 1). Five fresh flower pollen mocks (M0001-0005) were constructed. Among these, M0001 was made with an equal pollen ratio, which was used for calibrations of plastid genome copy numbers (See “Genome-skimming of Mock Pollen Mixtures”). Species ratios in FP mocks were set to test pollen-number variation from a minimum of 1-fold (e.g., *Al. aurea* vs. *L. brownii* in M0003) to a maximum of 100-fold (e.g., *L. brownii* vs. *Ab.* spp. in M0004). Fourteen bee pollen mocks were constructed, with M0014-0018 and 0021 (equal species ratio at varied total counts) used for testing repeatability of the proposed protocol and for calibrations of plastid genome copy numbers of the relevant bee pollen species. M0006 and M0007 were mock sample replicates, while M0008 and M0009 were DNA replicates. Species ratios in the rest of BP mocks (M0010-0013) were set to test pollen number variations from a minimum of 1-fold (e.g., *B. napus* vs. *Pa. rhoeas* in M0010) to a maximum of 300-fold (e.g., *C. japoica* vs. *Pr. armeniaca* in M0010).

**Table 1.** Total pollen counts and species ratios in mock samples.

|  | Mock samples | Total pollen counts | Species ratios |
| --- | --- | --- | --- |
|  |  |  | *L. brownii* : *Ab. pictum* : *Al. aurea* : *S. truncates* : *Ab. megapotamicum* : *An. majus* : *N. stellata* |
| FP | M0001 | 500,000 | 1 : 1 : 1 : 1 : 1 : 1 : 1 |
|  | M0002 | 500,000 | 91 : 27 : 9 : 3 : 1 : 9 : 1 |
|  | M0003 | 500,000 | 1 : 9 : 1 : 30 : 9 : 15 : 30 |
|  | M0004 | 500,000 | 10 : 1000 : 100 : 10 : 1 : 0 : 0 |
|  | M0005 | 500,000 | 4 : 16 : 32 : 2 : 1 : 2 : 8 |
|  |  |  | *R. chinensis* : *B. napus* : *Pa. rhoeas* : *V. faba* : *C. japoica* : *Pr. armeniaca* |
| BP | M0010 | 2,500,000 | 25 : 5 : 5 : 125 : 1 : 300 |
|  | M0011 | 2,500,000 | 180 : 90 : 30 : 30 : 1 : 1 |
|  | M0012 | 2,500,000 | 30 : 90 : 1 : 1 : 180 : 30 |
|  | M0013 | 2,500,000 | 4 : 16 : 2 : 32 : 1 : 8 |
|  | M0006 | 2,500,000 | 5 : 300 : 125 : 25 : 5 : 1 |
|  | M0007 | 2,500,000 | 5 : 300 : 125 : 25 : 5 : 1 |
|  | M0008 & M0009 | 5,000,000 | 5 : 300 : 125 : 25 : 5 : 1 |
|  | M0014 | 200,000 | 1 : 1 : 1 : 1 : 1 : 1 |
|  | M0015 | 500,000 | 1 : 1 : 1 : 1 : 1 : 1 |
|  | M0016 | 1,000,000 | 1 : 1 : 1 : 1 : 1 : 1 |
|  | M0017 | 2,500,000 | 1 : 1 : 1 : 1 : 1 : 1 |
|  | M0018 | 2,500,000 | 1 : 1 : 1 : 1 : 1 : 1 |
|  | M0021 | 5,000,000 | 1 : 1 : 1 : 1 : 1 : 1 |

M0001–0005 are FP mixtures; M0006–0018 and M0021 are BP mixtures. M0006 and M0007 are sample replicates; M0008 and M0009 are DNA replicates. M0014–0018 and M0021 contain pollen at equal ratios but varied total pollen counts.

### Pollen DNA Extractions

Pollen DNA was extracted using the Wizard method (Soares, Amaral, Oliveira, & Mafra, 2015), where the Wizard columns were replaced by Genomic Spin Columns (Transgen Biotech, Beijing, China).

We also estimated the number of pollen grains needed to produce the regular DNA mass required for the library construction on Illumina platforms (200 ng). The DNA yield per pollen grain is expected to vary among species due to differences in nuclear genome sizes and plastid genome copy numbers. Total DNA was extracted from 20,000, 40,000, 100,000, 200,000 pollen grains of *B. napus, C. japonica, Pa. rhoeas, Pr. armeniaca, Rh. chinensis*, *V. faba* and a mixture containing all 6 BP species at an equal ratio. DNA extraction was repeated 3 times for each species at each pollen count (12 extracts for each species), each of which was quantified using an Invitrogen Qubit®3.0 Fluorometero.

### Bee Pollen Barcoding

The taxonomic identifications of BP were confirmed by Sanger sequencing of the *rbcL* barcodes. Pollen DNA was extracted as described above. One microliter of each primer (1F: 5’- ATGTCACCACAAACAGAAAC-3’ and 724R: 5’-TCGCATGTACCTGCAGTAGC-3’, Fay, Bayer, Alverson, De Bruijn, & Chase, 1998) was used in a PCR reaction with a total volume of 20 μL, containing 2.5 μL of 10x *TransStart Taq* Buffer, 3.2 μL of dNTP (Promega U1515), 0.2 μL of *TransStart Taq* DNA Polymerase and 1 μL of template DNA. The PCR program was set as: initial denaturation at 95°C for 2 min, 34 cycles of 94°C denaturation for 1 min, annealing at 55°C for 30 s, and extension 72°C for 1 min, and a final extension step at 72°C for 7 min. Amplicons were sequenced using Sanger sequencing at Ruibiotech, Beijing, China. Sanger sequences were blasted individually against the GenBank nucleotide database for taxonomic identifications.

### Construction of a Reference Database for Plastid Genomes

About 100 mg of dried leaf tissues of each fresh flower species was ground with liquid nitrogen, treated with solution A following a modified CTAB method (Li, Wang, Yu, Wang, & Zhou, 2013) and then extracted using a Plant Genomic DNA Kit (TIANGEN, Beijing, China).

Libraries with an insert-size of 350 bp were prepared using leaf DNA extracts of *Ab. pictum*, *Ab. megapotamicum*, *An. majus*, *N. stellate* and *S. truncates*, following the manufacturer’s instruction. DNA libraries were sequenced at 2 Gb per species with 150 paired-end (PE) reads using an Illumina HiSeq4000 at BGI-Shenzhen, China. Additionally, the pollen DNA of *L. brownii* was sequenced with the same sequencing strategy using a HiSeq X Ten at NOVOgene (Beijing, China).

Data filtering removed reads containing adaptor contamination, duplication contamination, poly-Ns (>15 Ns) and those of >60 bases with quality score ≤32. Assemblies of chloroplast genomes were conducted using NOVOPlasty (Dierckxsens, Mardulyn, & Smits, 2017), except for *L. brownii*, which was assembled using SOAPdenovo-Trans (K=71, Xie et al., 2014). Protein-coding genes (PCGs) were annotated by using perl scripts from Zhou et al. (2013), which blasted the assemblies against a database containing 1,552 angiosperm chloroplast genomes (Table S2 in Appendix 1) downloaded from GenBank and predicted putative PCGs. All predicted plastid PCGs were aligned using MEGA 7.0 (Kumar, Stecher, & Tamura, 2016), and then PCGs shared by all FP and all BP taxa were concatenated respectively and used as respective reference sequences for pollen mixture analysis (Fig. 1).

**Fig. 1.**
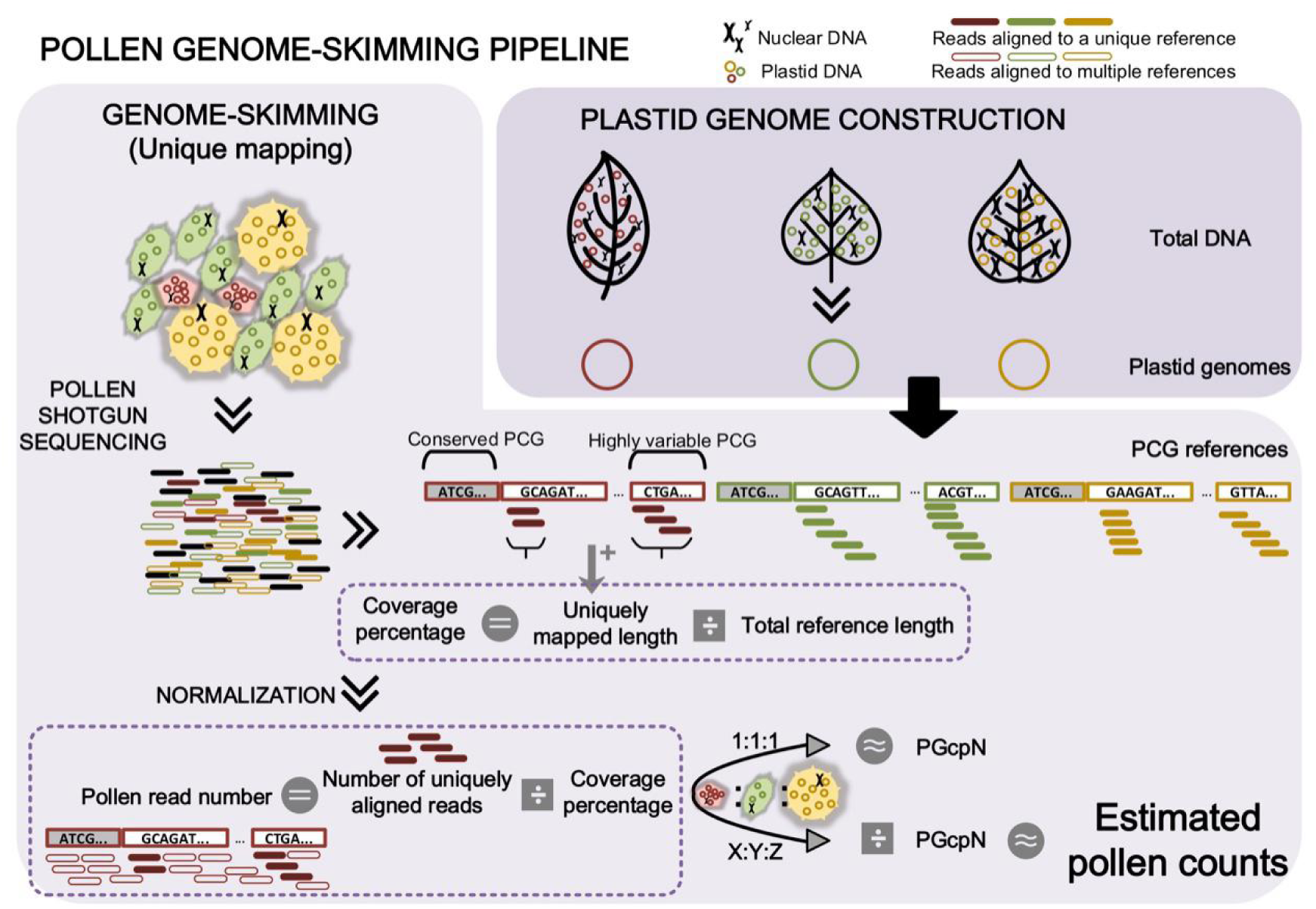
Pollen genome–skimming pipeline. Plastid protein–coding gene (PCG) references were obtained from “PLASTID GENOME CONSTRUCTION” and used for unique mapping in “GENOME– SKIMMING”. Only reads mapped onto a unique reference (solid short bars) were retained for calculation of the coverage percentage, which was then used for normalization of pollen read number for each member species. Pollen read numbers of member species in mock samples mixed at equal pollen counts (1:1:1) were considered as the plastid genome copy number (PGcpN) for corresponding species. The PGcpN was then used in calculating pollen counts for all member species from sequence reads in regular mocks (X:Y:Z).

### Genome-skimming of Mock Pollen Mixtures

For each mock, about 200 ng of DNA was used for library construction and high-throughput sequencing. A 350 bp insert-size library was sequenced at 4 Gb depth and 150 PE on an Illumina HiSeq4000 platform for each mock pollen mixture samples. After data filtering as described above, clean reads for BP and FP samples were mapped onto reference PCGs using *aln* BWA 0.7.16 (Li & Durbin, 2010). Aligned reads were assigned to the mapped species only if they met all following criteria: 100% read coverage, ≤1 base difference and aligned with no more than 1 reference (unique mapping). By the nature of the unique-mapping algorithm, uniquely mapped reads would only represent highly variable regions among reference genomes. Additional reads would be expected to match multiple PCGs of low taxonomic resolutions, which would not be assigned to any specific taxon (Tang et al., 2015). Therefore, the total pollen read number of a given species was defined as the number of uniquely mapped reads divided by the coverage percentage of its reference PCG sequence (Fig. 1).

The copy number of plastid genomes in matured pollen show drastic variations between plant species but remain relatively conservative within species (personal communication with Dr. Sodmergen of Peking University, China). Mock samples M0001 (FP), M0014-18 (BP) and M0021 (BP) were constructed with all member species mixed at equal pollen ratios. Therefore, in the sequencing results, proportions of sequence read of the member species are expected to reflect natural copy number differences in plastid genomes among species. These 7 mock samples were then used to estimate relative plastid genome copy number (PGcpN) ratios among BP and FP (Fig. 1). The PGcpN of the species with the least number was set as 1 and the average values of BP replicates (M0014-18 and M0021) were adopted as PGcpN ratios for the member species. For other pollen mixture samples, the pollen read number for each species per sample was then weighted by its PGcpN ratio, resulting in its estimated pollen count in the mixture (Fig. 1, Table S3 in Appendix 1). Linear regression between pollen proportion from Table 1 and pollen frequency computed from sequencing reads was performed in R 3.4.4 base package (R Core Team 2015), to estimate correlations between pollen counts and sequence reads for all species.

### Examination for Species Mixture in BP

As honeybees are generalist pollinators, each pollen pellet collected from the corbicula is expected to contain pollen from multiple plants. To examine to what extent the purchased BP are “contaminated” by non-labeled pollen species, we sequenced each BP at 2 Gb with 150 PE reads on Illumina platforms (*B. napus, R. chinensis* and *Pa. rhoeas* on a HiSeq X Ten at Novogene, Beijing, China; *Pr. armeniaca, C. japoica* and *V. faba* on a HiSeq 4000 at BGI-Shenzhen, China). Clean data were uniquely mapped onto BP reference PCGs as described above and used to compute the read percentage for each of the mixed species.

## Results

### DNA Extraction from Pollen Samples

A total of 900 - 1500 ng of DNA were obtained for the tested pollen mocks (Table S4). These results confirmed that 1 or 2 pollen pellets carried by a single honeybee can usually provide more than enough DNA for high-throughput sequencing.

DNA yields were positively correlated to the number of grains used for extraction within each species (Fig. 2, Table S5 in Appendix 1). On the other hand, species showed consistent differences in DNA yields. For instances, ca. 200,000 grains were needed to produce 200 ng DNA in *Pr. armeniaca*, whereas only ca. 40,000 grains needed for *C. japoica.* These differences are caused by variations in genome sizes and plastid genome numbers in pollen across species. Approximately 60,000 pollen grains of the mixed sample (consisting of 6 species at equal ratios) were required for 200 ng DNA.

**Fig. 2.**
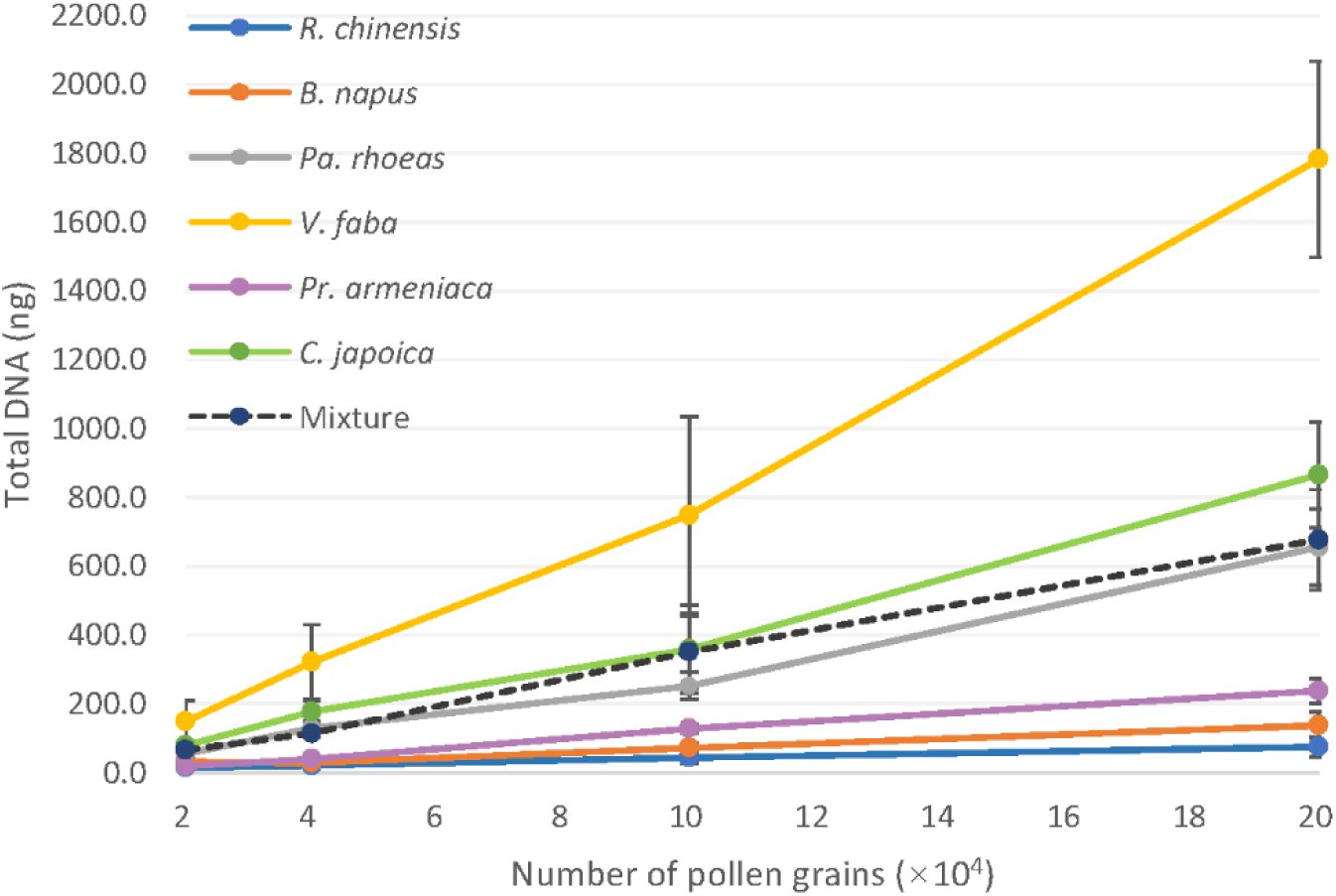
DNA yields from six bee pollen species. DNA were isolated respectively from 20,000, 40,000, 100,000 and 200,000 pollen grains of each BP species and the mixture contained six pollen species at equal ratios.

### **Bee Pollen Identification by** *rbcL* **Barcoding**

Sanger sequences of the *rbcL* barcodes for the six BP species were obtained (Table S6 in Appendix 1) with high quality. Barcodes confirmed the taxonomic identifications for the labeled species at ⩾99% identity.

### Plastid Reference Genome

Plastid genomes were assembled into scaffolds of 120-167 kb for five plant species (Fig. S1 in Appendix 2). Reads assigned to plastid genomes account for 2.8-19.6% of the total shotgun reads for varied species, with an exception in *L*. *brownii* that was extracted from pollen, which contained much fewer plastid genome copies than did leaf tissue (0.07%, Table S7 in Appendix 1). In addition, plastid genomes of *B. napus* (NC_016734.1), *C. japoica* (NC_036830.1), *R. chinensis* (NC_033535.1), *Pr. armeniaca* (KY420025.1), *Pa. rhoeas* (MF943221.1), *V. faba* (KF042344.1) and *Al. aurea* (KC968976.1) were downloaded from GenBank and were included in the reference. A total of 65 PCGs shared by all FP species and 71 shared by all BP species were used as reference PCGs, respectively. However, *Abutilon megapotamicum* and *Ab. pictum* could not be differentiated from each other even using 65 PCGs, due to limited taxonomic resolution of the gene markers. These two species were pooled for downstream analysis.

### Copy Number Variation of Plastid Genomes among Pollen Species

Mock samples were constructed with equal pollen counts for all member species, for both FP (M0001) and BP (M0014-0018, 0021). Shotgun reads were assigned to species using our unique-mapping criteria and pollen read number for each species per sample was calculated as described above. The species assigned with the least pollen read number was used as the standard, against which pollen read number of all other species were compared to produce relative copy numbers of plastid genomes (PGcpN). In our results, member species showed significant variations in plastid genome numbers (not the number of plastid organelles). For instance, *Al. aurea* had 27.8 times more plastid genomes per pollen than that of *An. majus*, and *Pr. armeniaca* had 57.7 times more plastid genomes per pollen than that of *B. napus* on average (Table 2). Nevertheless, plastid genome numbers were conserved within species as shown in BP samples M0014-0018 and M0021 (Table S3 in Appendix 1) and in proportion results in BP replicates M0006-0009 (see “Repeatability”).

**Table 2.** Pollen proportions of mock samples calculated from pollen counts and sequences.

|  |  | Pollen counts |  |  |  |  |  | Sequence reads |  |  |  |  |  |
| --- | --- | --- | --- | --- | --- | --- | --- | --- | --- | --- | --- | --- | --- |
|  |  | *L. brownii* | *Ab. spp.* | *Al. aurea* | *S. truncates* | *An. majus* | *N. stellata* | *L. brownii* | *Ab. spp.* | *Al. aurea* | *S. truncates* | *An. majus* | *N. stellata* |
| PGcpN |  |  |  |  |  |  |  | 3.4 | 2.2 | 27.8 | 21.1 | 1.0 | 9.5 |
| FP | M0002 | 64.5% | 19.9% | 6.4% | 2.1% | 6.4% | 0.7% | 23.7% | 63.1% | 2.1% | 1.3% | 8.9% | 0.9% |
|  | M0003 | 1.1% | 18.9% | 1.1% | 31.6% | 15.8% | 31.6% | 1.1% | 20.8% | 1.4% | 26.5% | 11.4% | 38.8% |
|  | M0004 | 0.9% | 89.3% | 8.9% | 0.9% | 0.0% | 0.0% | 1.0% | 91.1% | 5.5% | 0.6% | 1.4% | 0.4% |
|  | M0005 | 6.2% | 26.2% | 49.2% | 3.1% | 3.1% | 12.3% | 7.2% | 29.3% | 40.2% | 2.7% | 8.9% | 11.7% |
|  |  | *R. chinensis* | *B. napus* | *Pa. rhoeas* | *V. faba* | *C. japoica* | *Pr. armeniaca* | *R. chinensis* | *B. napus* | *Pa. rhoeas* | *V. faba* | *C. japoica* | *Pr. armeniaca* |
| PGcpN |  |  |  |  |  |  |  | 1.4 | 1.0 | 3.9 | 2.9 | 2.0 | 57.7 |
| BP | M0006 | 1.1% | 65.1% | 27.1% | 5.4% | 1.1% | 0.2% | 3.2% | 56.3% | 31.0% | 6.5% | 2.6% | 0.4% |
|  | M0007 | 1.1% | 65.1% | 27.1% | 5.4% | 1.1% | 0.2% | 2.9% | 60.1% | 28.0% | 6.6% | 2.1% | 0.3% |
|  | M0008 | 1.1% | 65.1% | 27.1% | 5.4% | 1.1% | 0.2% | 3.9% | 57.6% | 26.1% | 9.1% | 3.0% | 0.3% |
|  | M0009 | 1.1% | 65.1% | 27.1% | 5.4% | 1.1% | 0.2% | 3.6% | 55.8% | 27.6% | 9.9% | 2.8% | 0.3% |
|  | M0010 | 5.4% | 1.1% | 1.1% | 27.1% | 0.2% | 65.1% | 7.1% | 6.4% | 1.6% | 31.7% | 5.3% | 47.9% |
|  | M0011 | 54.2% | 27.1% | 9.0% | 9.0% | 0.3% | 0.3% | 57.4% | 21.4% | 10.0% | 9.1% | 1.7% | 0.4% |
|  | M0012 | 9.0% | 27.1% | 0.3% | 0.3% | 54.2% | 9.0% | 11.1% | 24.4% | 0.9% | 1.0% | 52.9% | 9.8% |
|  | M0013 | 6.3% | 25.4% | 3.2% | 50.8% | 1.6% | 12.7% | 7.7% | 25.0% | 3.4% | 49.4% | 3.8% | 10.7% |

M0006–0009 are BP mock samples used for repeatability test. Table cells were colored in red, with the darkest representing the highest percentage. The two red blocks were putative false positives detected by sequencing.

### Repeatability

The sequencing results were highly repeatable in both sets of BP mock samples constructed at different species ratios (Fig. 3, Table 2). It is worth noting that these results also reflect consistency in all relevant steps involved in the pipeline, including pollen counting, subsampling, pollen pooling, DNA extraction, library construction, sequencing and etc.

**Fig. 3.**
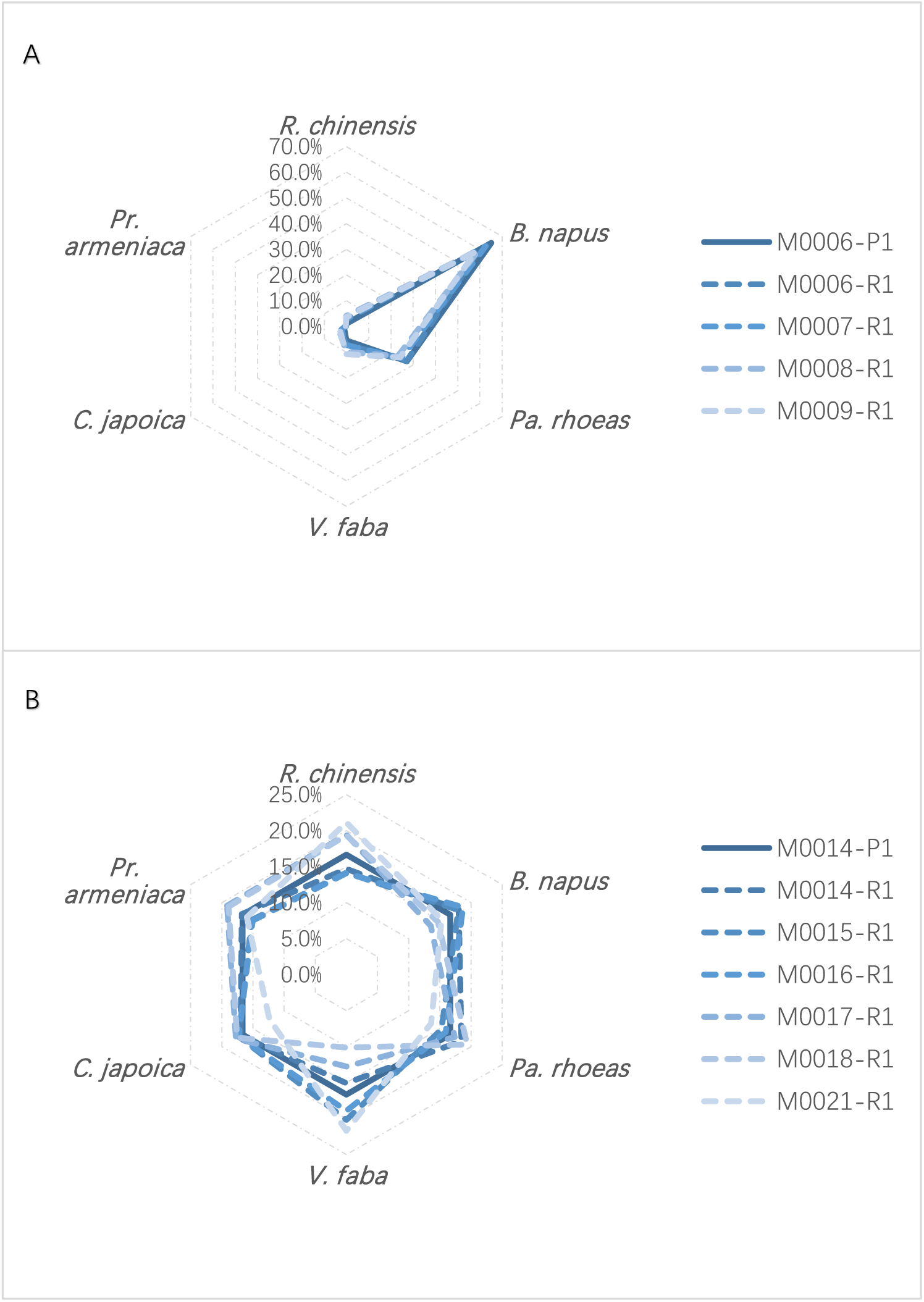
Repeatability of pollen genome–skimming. Mock samples consisted of 6 species of fresh flower pollen at given ratios: *R. chinensis: B. napus: Pa. rhoeas: V. faba: C. japoica: Pr. armeniaca* = 5:300:125:25:5:1 (panel A, 4 replicates) and 1:1:1:1:1:1 (panel B, 6 replicates). Solid lines and “–P1” represent results from pollen counts; dashed lines and “–R1” represent replicates of genome– skimming sequencing.

### Species Richness and Abundance of Pollen Mocks

Genome-skimming successfully detected all pollen species in all mock samples (Table 2, Table S3 in Appendix 1), including species found at just 0.2% of the total abundance (*C. japoica* in M0010, and *Pr. armeniaca* in M0006-0009). In principle, our stringent unique-mapping criteria would produce conservative results, where the reference species uniquely mapped by sequence reads would unlikely be an analytical artefact. This method was deliberately chosen to alleviate issues associated with the high sensitivity of high-throughput sequencing technologies, where they tended to pick up minute traces of DNA from the environment, causing false positives. Following this method, low-frequency species identified by sequencing are likely present in the real sample, providing confidence in detecting rare species in sample mixtures. In fact, all species with low abundances (e.g., with a relative abundance of 0.2% by pollen counting) were readily detected (Table 2, Table S3 in Appendix 1). However, two species absent from pollen counting were also detected by sequencing (*An. majus* and *N. stellata* in M0004). These 2 species were not pooled in the mock sample but showed non-negligible read depths and coverages (2.1X, 43.9% for *An. majus* and 5.1X, 23.2% for *N. stellata*, respectively), which were comparable to rare taxa truly present in M0002 (2.0X, 46.0% and 1.9X, 34.9% for the corresponding species, respectively) (Table S3 in Appendix 1). These results suggested that the two species detected by sequencing were likely a result of sample contamination rather than being analytical artefacts.

Quantification results for nearly all FP and BP mock samples were highly congruent with those from pollen counting (Table 2, Figs. 4, 5). The radar shapes representing species compositions concluded from sequencing (Fig. 3, dashed lines) matched well with those from pollen counting (Fig. 3, solid lines). The only exception was M0002, where the relative abundance (pollen ratio in mixture) of *L. brownii* was lower in the sequencing result than pollen counting, while that for *Abutilon* spp. was overestimated by the sequencing approach. This result was consistent in our repeat (Fig. S2 in Appendix 2, Table S3 in Appendix 1).

**Fig. 4.**
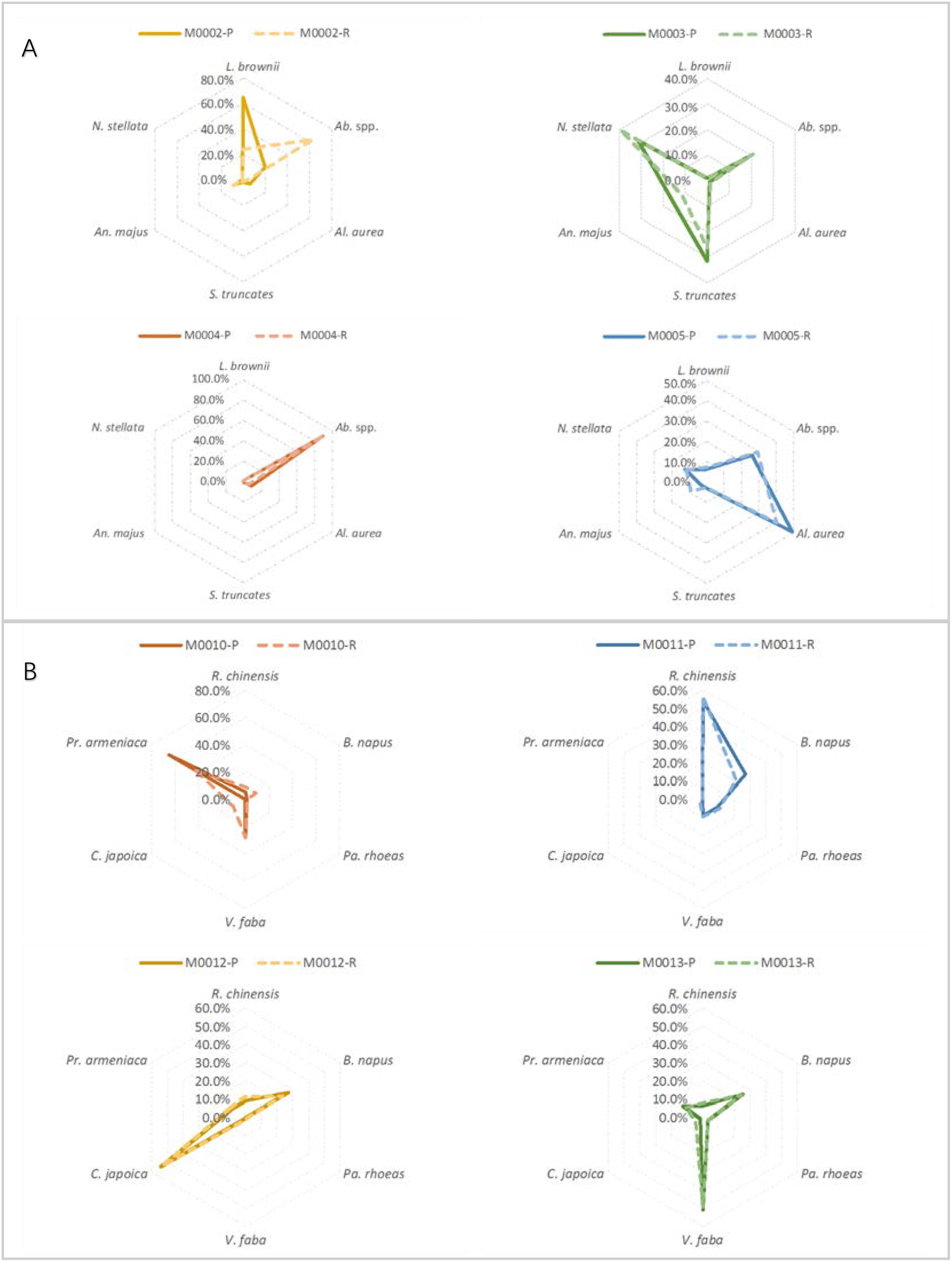
Coherence in pollen proportions between counting and sequencing. Solid lines represent results from pollen counting and dashed lines represent replicates of genome skimming sequencing.

**Fig. 5.**
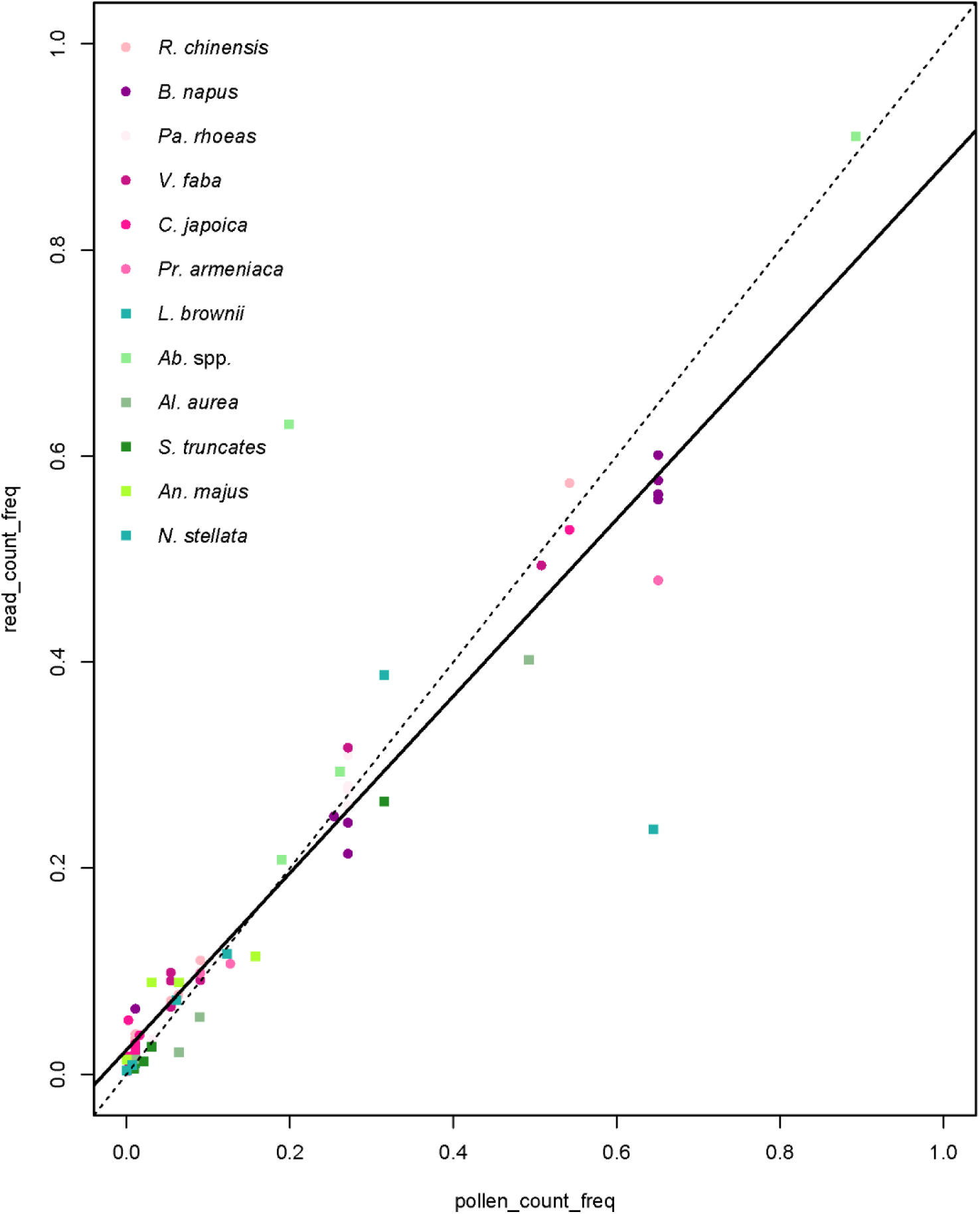
Scatter plots of pollen species frequencies from pollen counts versus genome– skimming. The solid line is the linear regression (read_count_freq ∼ 0.02367 + 0.85800*pollen_count_freq) and the dashed line is the 1:1 line, representing complete match between results from pollen counts and sequence reads.

Overall, pollen frequencies computed from pollen reads of each species were significantly correlated with corresponding pollen count proportions (linear model, R^2^ =86.7%, P = 2.2e-16, Fig. 6). The genome-skimming approach demonstrated a high level of sensitivity and accuracy in discriminating pollen counts at a wide range of compositional differences. In 68 out 70 cases (97.1%), the sequencing results were able to quantify pollen proportions to the correct order of magnitudes. Even when the maximum differences among pollen species had reached more than 100-fold within mock samples (e.g., M0004, 0006-0012), genome-skimming was still able to correctly quantify species at low abundances to the correct order of magnitudes with high consistency (e.g., *R. chinensis vs. C. japoica* in M0006-0009, *Pa. rhoeas* vs. *V. faba* in M0012).

### Pollen Mixture in Bee Pollen

The majority of each BP data set were mapped back onto the corresponding reference genomes, indicating that the bee pollen pellets were mostly made up of the labeled pollen species. However, it was also clear that all BP samples were mixed with other pollen species at varied proportions (Table 3). BP *V. faba* contained the most non-labeled pollen DNA (33.97%), while *Pr. armeniaca* showed the least mixture (1.16%). These results suggest that honeybees may regularly visit multiple flower species in a single trip, although we could not rule out the possibility of sample mixing during bee pollen preparation by the merchandise.

**Table 3.** Pollen mixing frequencies in bee pollen suggested by sequencing data.

| Reference<br>species<br>BP sample | *R. chinensis* | *B. napus* | *Pa. rhoeas* | *V. faba* | *C. japonica* | *Pr. armeniaca* |
| --- | --- | --- | --- | --- | --- | --- |
| *R. chinensis* | 84.28% | 2.68% | 0.00% | 0.00% | 7.21% | 5.83% |
| *B. napus* | 0.00% | 85.74% | 6.34% | 0.00% | 4.75% | 3.17% |
| *Pa. rhoeas* | 0.00% | 0.00% | 96.63% | 0.00% | 1.56% | 1.82% |
| *V. faba* | 5.47% | 5.93% | 4.06% | 66.03% | 9.94% | 8.56% |
| *C. japonica* | 2.98% | 3.27% | 2.96% | 2.80% | 82.52% | 5.47% |
| *Pr. armeniaca* | 0.37% | 0.15% | 0.07% | 0.17% | 0.41% | 98.84% |

All BP samples were primarily consisting of the labeled species, but with multiple sources of pollen mixing at varied proportions.

## Discussion

Pollination networks are complicated by nature, where visitation frequencies, pollen transport efficiencies, variations in pollen deposition and plant reproductive success have all been considered in various network construction methods. Recent studies suggested that these parameters would complement each other and collectively produce a better network (Ballantyne, Baldock, & Willmer, 2015). However, it is also apparent that balance would have to be made between an accurate construction of a comprehensive network and the overall efficiency in ecological studies, especially those at large scales. Therefore, an effective approach to identifying pollen diversity for flower visitors would help to incorporate this important information into the construction of pollination networks.

Although some studies have suggested metabarcoding is able to estimate valid abundances for pollen mixtures using amplicon frequencies (Pornon et al., 2016), others have shown less reliable correlations (Keller et al., 2015; Richardson et al., 2015). These conflicting observations may imply that the success of PCR-based metabarcoding is dependent on species composition of the pollen sample, which is highly variable in natural systems. By bypassing target gene amplifications and by expanding sequence references, the genome-skimming method can further provide quantitative pollen compositions for individual bees (corbicula pollen pellets) or pooled bees (pollen grains on bee bodies). In our results, a consistent positive correlation between pollen counts and pollen read numbers was established, in congruence with previous studies on macro-invertebrates. In fact, the correlation of the linear model is much more significant in pollen than in invertebrate animals tested so far (Tang et al., 2015; Bista et al., 2018), which is likely due to a higher level of homogeneity in organelle genome copy numbers per sample unit (pollen grain vs. individual animal). As with previously studied animal samples, this abundance-read correlation was significant independent of phylogenetic relationships among member pollen species (valid in both FP and BP samples) and levels of heterogeneity in pollen proportions (from 1-fold to 300-fold). The only exception in our study was observed in M0002, where the proportions of *L. brownii* and *Abutilon* spp. were seemingly flipped in the sequencing results. This result was repeated in our second trial (Fig. S2 in Appendix 2, Table S3 in Appendix 1), which excluded the likelihood of errors in sample contamination or mis-labeling. We speculate that the structural nature of the *Lilium* pollen (more hydrophobic compared to other pollen) may have caused its reduced abundance in final pollen mixtures, in which case the sequencing results would be more reliable. In fact, *Lilium* pollen floated in the supernatants, which might have led to its low representation in the mock subsamples.

Our pipeline also expanded reference gene markers from standard DNA barcodes (*matK* and *rbcL* for plants) to dozens of PCGs selected from whole plastid genomes. This extended sequence reference can produce better taxonomic resolution by recruiting additional variable genes, although some closely related species may still remain indistinguishable, as demonstrated by *Ab. megapotamicum* and *Ab. pictum* in our study. As with classic DNA barcoding approach, incomplete reference databases are often a key factor in causing false negatives. Fortunately, HTS-based methods have promised feasible paths in producing both standard DNA barcodes (Liu, Yang, Zhou, & Zhou, 2017; Hebert et al., 2018; Srivathsan et al., 2018) and organelle genomes (Straub et al., 2011; Tang et al., 2014) at significantly reduced costs. Indeed, large sequencing efforts for chloroplast genomes have seen significant progress in China. By November 2017, plastid genomes of 4,000 plants have been sequenced by the Kunming Institute of Botany, China. And an ambitious plan is in place, with a goal to complete the sequencing of plastid genomes for 18,000 Chinese seed plant species by 2021, covering ca. 2,750 genera (Li et al. in review).

While gaining benefits in producing quantitative results, the genome-skimming approach shows some compromise, where it requires higher DNA quantity for library construction and HTS sequencing (Zhou et al., 2013). Considering the low unit weight of pollen grains, DNA quantity may present a major challenge to a PCR-free based method. Current Illumina-based sequencing protocols require 200 ng or less genomic DNA for library construction, which is roughly the amount of DNA extracted from ca. 60,000 mixed pollen grains of tested species (Fig. 2). Regular pollen pellets collected from a single corbicula of the honeybees (*Apis mellifera*) were estimated to each contain more than 100,000 pollen grains (Table S1 in Appendix 1), which shall provide sufficient DNA for standard high-throughput sequencing. On the other hand, pollen carried on the body of pollinating bees may contain a much smaller number of grains and may vary significantly across species and individuals. Relevant studies are scarce, but some showed that honeybees and bumblebees visiting *Rhododendron ferrugineum* in the Alps carried ca. 11,000 and 15,000 pollen grains on the body of each insect, respectively (Escaravage & Wagner, 2004). In these cases, multiple individuals of bees (e.g., 10) would need to be pooled to reach the need for genome-skimming. It is worth noting that although the proposed method has a minimum requirement for the total DNA quantity, pollen species represented by low DNA proportions can still be readily detected from the mixture. For instance, our pipeline was highly sensitive in that all pollen species in the mocks were detected, including those with very low abundances (e.g., 0.2% for *C. japoica* in M0010, or ca. 100 pollen grains in a pollen mixture with ca. 47,000 total grains, Table S4 in Appendix 1).

Admittedly, most current pollen analysis employed in pollination network studies are based on individual pollinators. Modifications are needed to fully utilize the genome-skimming pipeline proposed in this study. We expect the following possibilities: 1) Further developments in high throughput sequencing technologies would continually reduce the minimum requirements for DNA quantity. In fact, DNA obtained from single cells has been proofed ample for genomics studies, provided with whole-genome amplifications (Wang & Navin, 2015). 2) Pollen pellets collected from bee corbiculae may contain similar pollen composition as those collected from their bodies, therefore providing reliable information for pollen transport web, with sufficient DNA for the genome-skimming method. Although some bees showed varied preferences in corbiculate pollen and body pollen through active or passive behaviors (Westerkamp, 1996), we expect that at least some bees would have more homogenous pollen preferences, especially generalist pollinators, which of course needs to be examined carefully. 3) New algorithm needs to be developed to incorporate heterogeneous pollination contributions for pollinators. Although specimen pooling (due to insufficient DNA) might mask variations among individuals, it also creates the potential to reduce stochastic errors, therefore providing better understanding on pollination diversity at the species or population level.

Analytical cost is obviously variable depending on service carriers and technology advents. At the time we conducted our study, the sequencing of *de novo* plastid genomes and pollen mixtures costed ca. 100 USD per sample (including DNA extraction, library construction, and sequencing at ca. 2-4 Gb data), with the prime on DNA library construction. Costs on both sequencing and library construction have seen significant reduction in the past decade, although the latter is at a much slower pace (Feng, Costa, & Edwards, 2018). Given current chemistry cost and the fact that most of the laboratory pipelines have been standardized and can be accomplished at regular molecular labs, we expect future cost on pollen genome-skimming can be further brought down to just a fraction of what we have now. Furthermore, global efforts on chloroplast genome sequencing coupled with more focused scrutiny of local flora will help to establish comprehend reference databases for molecular pollen identifications via metabarcoding or PCR-free genome-skimming.

Finally, in addition to applications in pollination networks, the proposed pollen genome-skimming method can also be useful in diet analysis of pollen consumers (bees, hoverflies, bee flies, beetles etc.), through sequencing gut contents. In these studies, DNA degradation may present a major challenge for a PCR-based method, but less so for a direct shotgun sequencing approach.

## Acknowledgements

We thank Dr. Lei Gu of Capital Normal University and Dr. Sodmergen of Peking University for their advice in plant taxonomy and knowledge on the biology and molecular mechanism of plant organelles. XZ acknowledge Dr. Douglass W. Yu of University of East Anglia, UK and Kunming Institute of Zoology, Chinese Academy of Sciences, for his helpful comments that improved the paper. XZ is supported by the National Natural Science Foundation of China (31772493) and funding from the Beijing Advanced Innovation Center for Food Nutrition and Human Health.

## Data accessibility

The genomic data sets (6 FP, 6 BP and 20 pollen mixture samples) have been deposited in GenBank (PRJNA481636).

## Author contributions

XZ, DDL, MT, and JHH designed the study. DDL and JHH conducted the bench work. DDL and MT performed data analysis. All authors participated in writing and proofed the manuscript.

